# Spontaneous allelic variant in *Ush1g* resulting in an expanded phenotype

**DOI:** 10.1101/2023.02.28.529432

**Authors:** Vladimir Vartanian, Jocelyn F. Krey, Paroma Chatterjee, Sherri M. Jones, Allison Curtis, Renee Ryals, R. Stephen Lloyd, Peter G. Barr-Gillespie

**Author notes:** Co-senior authors (R. Stephen Lloyd, Peter G. Barr-Gillespie).

## Abstract

Strategies to reveal the discovery of the relationships between novel phenotypic behaviors and specific genetic alterations can be achieved via either target-specific, directed mutagenesis or phenotypic selection following random chemical mutagenesis. As an alternative approach, one can exploit deficiencies in DNA repair pathways that are responsible for the maintenance of genetic integrity in response to spontaneously-induced damage. In the genetic background of mice deficient in the DNA glycosylase NEIL1, elevated numbers of spontaneous mutations arise from translesion DNA synthesis past unrepaired, oxidatively-induced base damage. Several litters of *Neil1* knockout mice included animals that were distinguished by their backwards-walking behavior in open-field environments, while maintaining frantic forward movements in their home cage environment. Other phenotypic manifestations included swim test failures, head tilting, and circling. Mapping of the mutation that conferred these behaviors revealed the introduction of a stop codon at amino acid 4 of the *Ush1g* gene; the allele was *Ush1g^bw^*, reflecting the backwards-walking phenotype. *Ush1g^bw/bw^* null mice displayed auditory and vestibular defects that are commonly seen with mutations affecting inner-ear hair-cell function, including a complete lack of auditory brainstem responses and vestibular-evoked potentials. As in other Usher syndrome type I mutant mouse lines, hair-cell phenotypes included disorganized and split hair bundles, as well as altered distribution of proteins for stereocilia that localize to the tips of row 1 or row 2. Disruption to the bundle and kinocilium displacement suggested that USH1G is essential for forming the hair cell’s kinocilial links. Due to the vestibular dysfunction, however, visual behavior as measured with optokinetic tracking could not be assessed in *Ush1g^bw/bw^* mice. Consistent with other Usher type 1 models, however, *Ush1g^bw/bw^* mice had no substantial retinal degeneration compared to *Ush1g*^*bw*/+^ controls out to six months. In contrast to previously-described *Ush1g* alleles, this new allele provides the first knockout model for this gene.

## Introduction

Strategies to identify and establish the functional significance of genes and their products has been pioneered through manipulation of genomes via increasingly complex experimental designs^1^. Many of these advances have been reduced to practice using the mouse genome as an experimental and screening platform. Insights gained into establishing the functional significance of individual genes has provided both the foundation from which to understand specific structure-function relationships and the framework to dissect complex pathways and processes. Although not a perfect surrogate for analyses of human genes or therapeutic design and validation, the mouse genome shares an 80% synteny with the organization of the human genome and a high percentage of readily identifiable orthologs. Thus, analyses of the phenotypic consequences of genotypic manipulation in the mouse serves as a strong foundational point from which to understand human disease^1^. The International Mouse Genotyping Consortium, whose primary objective is to coordinate the cataloging, design, production, and documentation of all human gene orthologs found within the mouse genome is close to achieving this goal, with documentation of nearly 17,000 gene knockouts and covering nearly 80% of the target goal^2^.

Along this discovery pathway, significant advances were made via the observation and collection of mouse phenotypes arising from not only spontaneous mutations, but also large-scale, random mutagenesis induced by exposures to ionizing radiation and chemical mutagenesis^1^. For strategies organized around chemical mutagenesis, several large international consortia utilized DNA alkylating agents such as ethylnitrosourea and ethylmethanesulfonate to create widespread base damage, located primarily on purines. The expectation was that these base modifications would ultimately lead to randomized mutagenesis of the entire genome. However, to mitigate the potentially deleterious effects of these base damages, cells possess a variety of DNA repair pathways, with small molecule DNA alkylation damage primarily removed by the base excision repair pathway (BER). BER consists of a series of coordinated sequential steps that are initiated by DNA glycosylases, which scan the genome for altered bases, flip the damaged nucleotide into a catalytic pocket, and initiate chemistry for base release via glycosyl bond cleavage, with the potential cleavage of the phosphodiester backbone. Following base release, the remainder of the pathway is devoted to the creation and filling in of short gaps in the strand from which the base was released followed by sealing the nick^3^. This is a generally high-fidelity pathway that relies on an undamaged complementary strand as a template for high-fidelity gap filling. However, if the base damage is not removed prior to DNA replication by high-fidelity DNA polymerases (α, δ and ε), blocked replication can be rescued following translesion DNA synthesis that is catalyzed by low-fidelity, low-processivity DNA polymerases^4^. This damage-tolerance mechanism is the primary mechanism for introducing point mutations and short deletions and insertions.

Thus, random chemical mutagenesis strategies to create saturation mutagenesis for phenotypic selection have relied on overwhelming DNA-repair pathways without inducing cytotoxicity. To the best of our knowledge, mice deficient in DNA-repair mechanisms have not been used in conjunction with large-scale chemical mutagenesis and subsequent selection. However, it would be anticipated that deficiencies in repair of spontaneously-induced base damage, including depurination, deamination, alkylation, and oxidation, as well as damage produced by exogenous sources, would continuously generate point mutations and short deletions. In this regard, base modification by exposure to reactive oxygen species is not only a by-product of normal biochemical reactions, but also from ionizing radiation, ultraviolet irradiation, heavy metal exposures, and inflammation^5^. Repair of the plethora of oxidatively-induced damages is initiated by a series of DNA glycosylases including NEIL1, NEIL2, NEIL3, NTH1, and OGG1 (ref. 6). NEIL1 is responsible for the removal of several saturated pyrimidines, both ring-fragmented purines, alkylated ring-fragmented purines, psoralen-induced DNA crosslinks, and secondary oxidation products of 8-oxoGua (ref. 7). Given the very broad substrate specificity of NEIL1, it would be anticipated that mice which are deficient in *Neil1* would show elevated levels of spontaneous mutagenesis. For over 20 years, the Lloyd laboratory has maintained *Neil^-/-^* mice, which are frequently bred as knockouts for metabolic and carcinogenesis studies. As part of routine colony expansion, mice are systematically monitored for unusual phenotypes. The current investigation arose by the observation of mice exhibiting a backwards-walking phenotype in an open-field environment, as well as exhibiting a variety of other abnormal behaviors. Since literature reviews did not yield matches to a backwards-walking phenotype, studies were initiated to identify the genetic alteration that produced this phenotype.

We mapped the backwards-walking trait to the *Ush1g* gene, and found that it was caused by a nonsense mutation that truncated the protein after the third amino acid. We found that *Ush1g^bw/bw^* mice had auditory and vestibular defects, as well as disruption of the sensory hair bundle of cochlear hair cells, much like that seen with other mouse lines mutant for Usher syndrome type I (USH1) genes. Examination of protein distribution by immunocytochemistry revealed that loss of *Ush1g* affects row 1 and row 2 tip proteins similarly to loss of *Cdh23* or *Pcdh15*. We noted defects in kinocilial location that support the hypothesis that USH1G assists in stabilizing CDH23-PCDH15 kinocilial links. Finally, like other mouse USH1 models, we detected no visual phenotypes in these mice. We conclude that *Ush1g^bw^* is a null *Ush1g* allele that will be useful for investigation of auditory and vestibular function.

## Materials & Methods

### Animal models

All animal procedures were approved by the Institutional Animal Care and Use Committee (IACUC) at Oregon Health & Science University (IP00000145 for RSL, IP00000610 for RR, and IP00000714 for PGBG). *Neil1^-/-^* mice have been described previously^8,9^. Mice arising from the original mutagenic event creating *Ush1g^bw^* were backcrossed to C57BL/6J (B6) mice (RRID:IMSR_JAX:000664, Jackson Laboratories, Bar Harbor, ME) and were maintained on a B6 background. Mouse pups were assumed to be born at midnight, so the animal age on the first day is referred to as P0.5. Both female and male pups were used for all experiments.

### Statistical analysis

Unless otherwise stated, statistical comparisons between two sets of data used the Student’s t-test with unpaired data and the untested assumption of equal variance. In figures, asterisks indicate: *, p < 0.05; **, p < 0.01; ***, p < 0.001.

### Optokinetic tracking

Optokinetic tracking (OKT) thresholds were used to identify spatial frequencies of gratings (cycles/degree), which define visual performance for animals (OptoMotry; CerebralMechanics, Lethbridge, Alberta, Canada). Briefly, animals were placed on the pedestal in the OptoMotry system and given five minutes to acclimate to the new environment. A simple staircase method at 100% contrast in normal lighting conditions was used for testing. Right and left eyes were tested separately and averaged together to get one spatial frequency per animal.

### Electroretinography

Mice were dark-adapted overnight. Under dim red light, mice were anesthetized with ketamine (100 mg/kg) and xylazine (10 mg/kg). Mouse pupils were dilated with 1% tropicamide and 2.5% phenylephrine and lubricated during the procedure with hypromellose 2.5% ophthalmic lubricant. Bilateral platinum electrodes were placed on the corneal surface to record the light-induced electroretinography (ERG) potentials. The reference and ground electrodes placed subcutaneously in the forehead and tail, respectively. Flashes with increasing light intensity from −4 to 3.39 log cd•s/m^2^ were used for the recordings. A 2-way ANOVA with Šídák’s multiple comparisons test was used to compare amplitudes between the groups.

### Retinal spectral domain-optical coherence tomography

At P180, mouse retinas were imaged with a Heidelberg Spectralis Multimodal Imaging Platform (Heidelberg Engineering Inc., Franklin, MA, USA) using spectral domain-optical coherence tomography (SD-OCT). Images were segmented using the Heidelberg Eye Explorer software included with the imaging platform. To measure photoreceptor thickness, REC+ was defined as the thickness from the base of the RPE to the interface of the inner nuclear layer and outer plexiform layer. REC+ values were taken from ten locations spanning superior, inferior, nasal and temporal retina. REC+ values from both eyes were averaged for each animal. A one-way ANOVA was used to compare photoreceptor thickness between groups.

### Retina histology

At P180, enucleated eyes were placed immediately in 4% formaldehyde and incubated overnight at 4°C. Eyes were processed and embedded in paraffin for sectioning (Tissue-Tek VIP 6, Tissue-Tek TEC 5; Sakura Finetek USA, Inc., Torrance, CA, USA). Sections were cut with a microtome to a thickness of 4 μm, stained with hematoxylin-eosin, and viewed on a Leica DMI3000 B microscope (Leica Microsystems GmbH, Wetzlar, Germany). A total of 12 images per eye encompassing the nasal, temporal, superior and inferior retinal quadrants were captured with a 40x objective. The number of photoreceptor nuclei in a row were counted and averaged from these 12 images, to obtain an outer nuclear layer (ONL) cell count per animal. A one-way ANOVA was used to compare ONL cell counts between groups.

### Retina immunofluorescence

At P180, WT, Het, and KO eyes were harvested and placed in 4% formaldehyde. At 24 hours after fixation, whole eyes were placed in a 30% sucrose solution for 4 hours, and then frozen in Scigen Tissue-Plus O.C.T. Compound (Thermo Fisher Scientific, Waltham, MA, USA) at −78°C. Eyes were sectioned at 12 μm on a Leica CM1860 cryostat. Cross-sections were incubated at 4°C in primary antibody consisting of anti-rhodopsin (1D4) (1:200, Cat# NBP1-30046; RRID:AB_1968611; Novus Biologicals, Centennial, CO, USA), 0.3% Triton X-100, and 1% bovine serum albumin overnight. Tissues were rinsed and incubated in secondary antibody containing goat anti-rabbit Alexa 488 (1:500, Thermo Fisher Scientific Cat# A11008; RRID:AB_143165), 1% bovine serum albumin, and PBS (1X) for two hours at 20 °C. After incubation in secondary antibody, retinas were rinsed again and placed in a DAPI solution (1:2400, Thermo Fisher Scientific Cat# D1306) for 5 minutes. Retinas were imaged with a white light laser confocal microscope (Leica Microsystems TCS SP8 X). Z-stacks (spanning 14-23 μm in 1 μm intervals) were collected using a 63x objective. Confocal images were processed using the maximum intensity tool in ImageJ software (version 1.49; National Institutes of Health, Bethesda, MD, USA).

### Functional tests for hearing and balance

Auditory brainstem response (ABR) experiments were carried out as described previously^10^ with 7 *Ush1g*^*bw*^/+ animals and 7 *Ush1g^bw/bw^* animals. Animals were anesthetized with xylazine (10 mg/kg, i.m.; Animal Health Inc., Greeley, CO, USA) and ketamine (40 mg/kg, i.m.; Hospira, Inc., Lake Forest, IL, USA), and placed on a heating pad in a sound-isolated chamber. Needle electrodes were placed subcutaneously near the test ear, both at the vertex and at the shoulder of the test ear side. A closed-tube sound-delivery system, sealed into the ear canal, was used to stimulate each ear. ABR measurements used tone bursts with a 1 ms rise time, applied at 4, 8, 16, 24, and 32 kHz. Responses were obtained for each ear, and the tone-burst stimulus intensity was increased in steps of 5 dB. The threshold was defined as an evoked response of 0.2 μV from the electrodes.

Vestibular evoked potential (VsEP) experiments were carried out as described previously^11^ with 7 *Ush1g^bw^*/+ animals and 7 *Ush1g^bw/bw^* animals. Mice were anesthetized by intraperitoneal injection of ketamine and xylazine (18 and 2 mg/ml; 5-7 μl/g body weight), followed by maintenance doses as needed to maintain adequate anesthesia. Core body temperature was maintained at 37.0 ± 0.2°C using a homeothermic heating pad (FHC Inc., Bowdoin, ME, USA). For VsEP testing, linear acceleration ramps producing rectangular jerk pulses were generated and controlled using a National Instruments (Austin, TX, USA) data acquisition system and custom software. Mice were placed supine on a stationary platform and the head was secured within a spring clip coupled to a voltage-controlled mechanical shaker (Model 132-2; Labworks, Costa Mesa, CA, USA). The head was oriented with nose up and linear translation stimuli were presented in the naso-occipital axis parallel to the Earth-vertical axis. Vestibular stimuli consisted of 2 ms linear jerk pulses, delivered to the head using two stimulus polarities—normal, with an initial upward jerk, and inverted, with an initial downward jerk—at a rate of 17 pulses per second. Stimulus amplitudes ranged from +6 dB to −18 dB re: 1.0 g/ms (where 1 g = 9.8 m/s^2^), adjusted in 3 dB steps. A broadband forward masker (50-50,000 Hz, 94 dB SPL) was presented during VsEP measurements to confirm the absence of auditory components^12^. Signal averaging was used to extract the VsEP responses from the background electrophysiological activity. Ongoing electroencephalographic (EEG) activity was amplified (200,000x), filtered (300-3000 Hz, −6 dB points), and digitized beginning at the onset of each jerk stimulus (1024 points, 10 μs/point) to produce one primary response trace. For each stimulus intensity and polarity, 128 primary responses were averaged to produce an averaged response waveform. Four averaged response waveforms were recorded for each stimulus intensity (two waveforms recorded for normal stimulus polarity and two for inverted polarity). Final individual response traces were produced by summing one averaged response to each stimulus polarity and dividing the result by two, thus producing two response traces for each stimulus intensity for each animal.

### Scanning electron microscopy

Periotic bones with cochleas were dissected in Leibovitz’s L-15 medium (Thermo Fisher Scientific) from P8.5 or P21.5 littermates from *Ush1g^bw^* crosses. After isolating the periotic bone, several small holes were made to provide access for fixative solutions; encapsulated cochleas were fixed for an hour in 2.5% glutaraldehyde in 0.1 M cacodylate buffer supplemented with 2 mM CaCl_2_. Next, cochleas were washed with distilled water and the cochlear sensory epithelium was dissected out; the tectorial membrane was manually removed. The cochlear tissues were then transferred to scintillation vials and dehydrated in a series of ethanol and critical-point dried using liquid CO_2_. Samples were immobilized on aluminum specimen holders using a carbon tape and sputter coated with 3-4 nm of platinum. Samples were imaged using the Helios scanning electron microscope.

### Immunofluorescence sample preparation for cochlea

Inner ears from *Ush1g^bw^* littermates were dissected at the indicated ages in cold Hank’s balanced salt solution (Thermo Fisher Scientific Cat# 14025076), supplemented with 5 mM HEPES, pH 7.4 (dissection buffer). Small openings were made within the periotic bones to allow perfusion of the fixative. Cochleas were fixed in 4% formaldehyde (Cat# 1570; Electron Microscopy Sciences, Hatfield, PA, USA) in dissection buffer for 20-60 min at room temperature. In experiments using row 2 protein antibodies (EPS8L2, CAPZB), fixation was for 20 min; row 1 protein antibodies were not sensitive to fixative duration and were usually fixed for 60 min. Ears were washed in PBS, then cochleas were dissected out from the periotic bone and the lateral wall was removed. Cochleas were permeabilized in 0.2% Triton X-100 diluted in 1x PBS for 10 min and blocked in 5% normal donkey serum (Cat# 017-000-121; Jackson ImmunoResearch, West Grove, PA, USA) for 1 hour at room temperature. For staining against USH1C, cochleas were permeabilized and blocked for one hour in 10% normal donkey serum (Cat# 017-000-121; Jackson ImmunoResearch, West Grove, PA, USA) and 0.2% Triton X-100 diluted in 1x PBS. Organs were incubated overnight at 4°C with primary antibodies in blocking buffer (5% normal donkey serum diluted in 1x PBS) and then washed three times in 1x PBS. Dilutions were 1:250 for anti-acetylated tubulin; 1:250 for anti-GPSM2, 1:500 for anti-GNAI3; 1:250 for anti-EPS8, 1:200 for USH1C; 1:250 for anti-EPS8L2, 1:250 for anti-CAPZB. Tissue was then incubated with secondary antibodies, which were 2 μg/ml donkey anti-rabbit Alexa Fluor 488 (Thermo Fisher Scientific Cat# A21206; RRID:AB_2535792) and 2 μg/ml donkey anti-mouse Alexa Fluor 568 (Thermo Fisher Scientific Cat# A10037; RRID:AB_2534013); 1 U/ml CF405 phalloidin (Cat# 00034; Biotium, Fremont, CA, USA) was also included for the 3-4 hr room treatment. Cochleas were washed three times in PBS and mounted on a glass slides in ~50 μl of Vectashield and covered with a #1.5 thickness 22 x 22 mm cover glass (Cat# 2850-22, Corning) or a #1.5 thickness 22 x 40 mm cover glass (Thermo Fisher Scientific Cat# CLS-1764-2240).

Primary antibodies were as follows. Mouse anti-acetylated tubulin (clone 6-11B-1) was Cat# T6793 (RRID:AB_477585) from Sigma-Aldrich (St. Louis, MO). Rabbit anti-GPSM2 Cat# HPA007327 (RRID: AB_1849941) from Sigma-Aldrich. Rabbit anti-GNAI3, directed against a C-terminal mouse GNAI3 peptide (KNNLKECGLY), was Cat# G4040 (RRID: AB_259896) from Sigma-Aldrich. Mouse monoclonal anti-EPS8 (clone 15), against mouse EPS8 amino acids 628-821, was Cat# 610143 (RRID: AB_397544) from BD Bioscience (San Jose, CA). Mouse monoclonal anti-CAPZB2, against a C-terminal peptide of human CAPZB2, was Cat# AB6017 (RRID: AB_10806205) from EMD Millipore (Burlington, MA). Mouse monoclonal anti-EPS8L2, against human EPS8L2 amino acids 615-715, was Cat# ab57571 (RRID: AB_941458) from Abcam (Cambridge, UK). Rabbit anti-pan-MYO15A was antibody PB48 from the Thomas Friedman lab^13^. We used Genemed Biosynthesis (San Antonio, TX) to generate a rabbit anti-USH1C antiserum using recombinantly expressed PDZ3 of mouse *Ush1c*. For attempted immunolocalization of USH1G, we used anti-sans J63 rabbit 73 (a gift from Amel El Bahloul-Jaziri).

### Fluorescence microscopy of stereocilia proteins

Organs were imaged using a 63x, 1.4 NA Plan-Apochromat objective on a Zeiss (Oberkochen, Germany) Elyra PS.1/LSM710 system equipped with an Airyscan detector and ZEN 2012 (black edition, 64-bit software) acquisition software. Settings for x-y pixel resolution, z-spacing, as well as pinhole diameter and grid selection, were set according to software-suggested settings for optimal Nyquist-based resolution. Raw data processing for Airyscan-acquired images was performed using manufacturer-implemented automated settings. Settings for x-y pixel resolution, z-spacing, as well as pinhole diameter and grid selection, were set according to software-suggested settings for optimal Nyquist-based resolution. Raw data processing for Airyscan-acquired images was performed using manufacturer-implemented automated settings. Display adjustments in brightness and contrast and reslices and/or maximum intensity Z-projections were made in Fiji software (https://imagej.net/software/fiji/).

## Results

### Discovery and identification of *Ush1g^bw^* mutant allele

As part of ongoing investigations into the role of the NEIL1 DNA glycosylase, routine expansion and health monitoring of the *Neil1^-/-^* colony revealed an unexpected phenotype. Because NEIL1 plays a major role in the initiation of base-excision repair of oxidatively-induced DNA damage and alkylated ring-fragmented purines^7^, throughout the maintenance of this strain, spontaneous mutations arise. Offspring of *Neil1^-/-^* crosses have generated mice with a variety of readily observable phenotypes, including but not limited to the generation of petite mice (<20 g at maturity), alopecia, microphthalmia, staggering, and seizures (unpublished observations).

In the current study and part of routine colony expansion of phenotypically normal *Neil1^-/-^* mice^9^, we observed a hyperactive male mouse. This mouse was mated with *Neil1^-/-^* females, generating phenotypically normal F1 offspring. When these offspring were set up in pair-wise mating, a portion of this F2 generation, representing both males and females, exhibited hyperactivity, head-tilting, and unresponsiveness to auditory stimuli. Further observation of these mice revealed that when picked up by the tail, they displayed tremors and rapid spinning. If they were placed in an open space context, such as a hood or a tabletop, they immediately stopped all forward motion and only walked backwards (Supplemental Video 1). Backward walking was also associated with backward circling, raising their heads to a vertical position, and head wagging. The affected mice could not perceive the edge of a surface and would routinely fall off the table top, mandating vigilance during behavioral monitoring. In their home cage, affected mice only walk or run forward and boxes containing only affected mice show rapid and sustained forward running in circular patterns within the cage. Affected mice were also unable to function in a swim test, rapidly swimming to the bottom of the tank and requiring rescue.

Subsequent mating in which both genders were phenotypically affected, or mating that used an affected female with a phenotypically normal male, resulted in successful pregnancies, albeit with small litters. All pups died within days after birth, however, presumably due to maternal disinterest or from being trampled during frantic, in-cage running. Analyses of mice generated from the mating of affected males with wild-type C57BL/6J females, followed by subsequent generation of F2 pups, revealed that the acquisition of these phenotypes was independent of the *Neil1* genotype.

To map the mutation, affected C57BL/6 males were mated with BALB/c females; the F1 generation mice were all phenotypically normal (Fig. 1). Offspring resulting from matings within this F1 generation produced a total of nine open-field, backwards-walking mice out of a total of 37 pups. Tails from a total of these nine mice and two additional backwards-walking mice and nine phenotypically normal mice were sent to The Jackson Laboratory for differential single nucleotide polymorphic (SNP) analyses. In this analysis, the approximate chromosomal location of the mutated gene was anticipated to map to inherited regions that were homozygous for C57BL/6 SNPs in the 11 affected mice and either heterozygotic or homozygotic for BALB/c SNPs in unaffected mice. Analyses of a total of 126 SNPs for each mouse revealed a single chromosomal location, extreme distal 11q that mapped to the mutated site (Fig. 1). The linkage was evident from SNP #11-117818566-N, marker RS3023766 at 116,784,04 bp. The boundaries of the gene of interest were established by SNP #11-105306632-N, marker RS3024066 at 104,384,279 bp and the end of chromosome 11 as the lower limit. Genes mapping to this general region were analyzed for potential functional interest, with *Ush1g* being the leading candidate. Sequence analyses revealed that the DNA encoding codon 4 of *Ush1g* (CAG, encoding glutamine) had been mutated to a stop codon (TAG). The mutant USH1G protein thus should be truncated after the third amino acid. Because of the previously-unreported backwards-walking phenotype of this allele, we designated it as *Ush1g^bw^* (MGI:7432804). Given the location of the introduced stop codon, we presume that the allele is a complete null.

**Figure 1.**
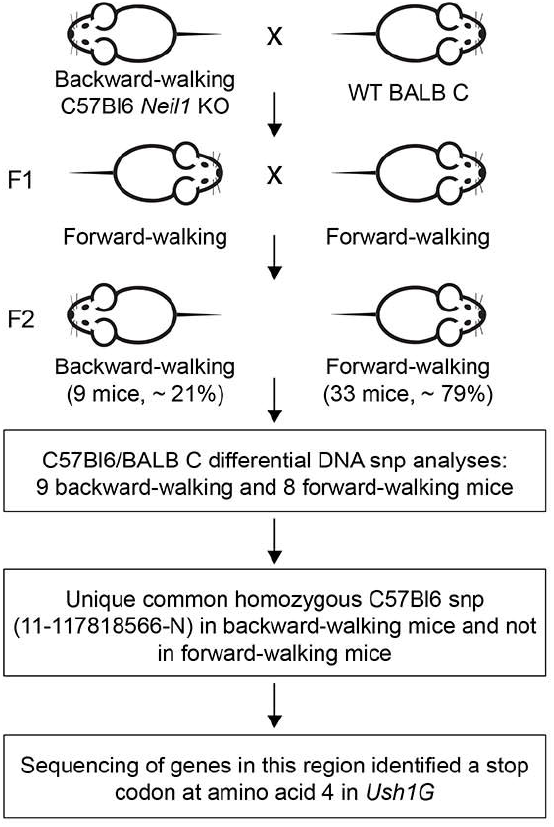
Scheme for identification of gene responsible for backwards walking. The mapping scheme used to identify *Ush1g^bw^* is shown. A backwards-walking *Neil1^-/-^* mouse was mated with a female BALB/c mouse; the F1 generation (all phenotypically wild type) was intercrossed to generate F2 mice. Roughly a quarter of the F2 generation had the backwards-walking phenotype. SNP analysis showed that SNP #11-117818566 was present in backwards-walking mice but not their phenotypically normal littermates. A nonsense mutation was identified in the *Ush1g* gene that terminates the protein after the third amino acid.

### Visual phenotype

*USH1G* causes progressive retinal degeneration in humans^14^. In *Ush1g^bw^* mice, visual behavior was measured in normal lighting conditions at P30, P90, and P180 using OKT; there were no significant differences between wild type and heterozygous *Ush1g* animals at these time points (Fig. 2A). By contrast, these measurements highlighted the *Ush1g^bw/bw^* phenotype; these animals were unable to balance on the platform, and measurements could not be acquired (Fig. 2A). Scotopic ERGs were recorded at these time points; while *Ush1g^bw/bw^* mice had significantly lower b-wave amplitudes than *Ush1g*^*bw*^/+ controls (Fig. 2C), a-wave amplitudes (corresponding to photoreceptor function) were essentially unchanged between the genotypes (Fig. 2B). At P180, prior to eye harvest for histology and immunofluorescence, SD-OCT was used to collect *in vivo* retinal images (Fig. 2D). Photoreceptor thickness calculated from SD-OCT images showed no difference between *Ush1g+/+, Ush1g^bw^/+* and *Ush1g^bw/bw^* animals (Fig. 2E). These results were confirmed with photoreceptor nuclei counts from histology images (Fig. 2F-G). Immunofluorescence of eyes harvested at P180 showed that rhodopsin levels in the photoreceptor outer segments were consistent between groups (Fig. 2H).

**Figure 2.**
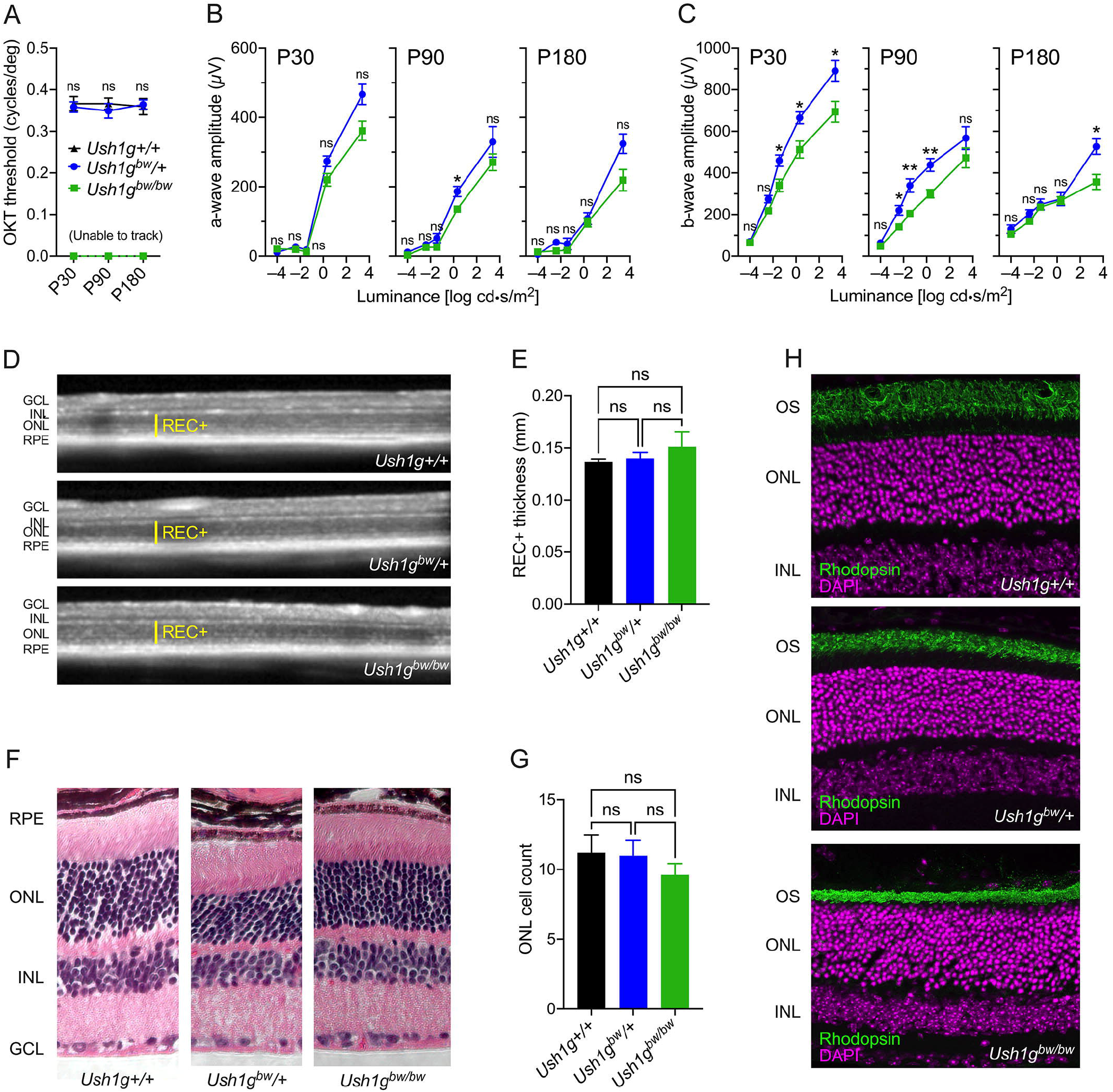
Retinal phenotype associated with the *Ush1g^bw^* mice. ***A***, Visual performance measured by optokinetic tracking (OKT). ***B-C***, Amplitudes of a-wave (B) and b-wave (C) generated from scotopic ERGs at range of light intensities. In each case, the three panels show results at P30, P90, and P180. Legend in A applies to panels B and C as well. Two-way ANOVA analyses were used to compare heterozygote and homozygote responses. ***D***, Representative SD-OCT images of the retina from *Ush1g+/+, Ush1g^bw^/+*, and *Ush1g^bw/bw^* mice. Yellow lines indicate photoreceptor layer thickness (REC+). ***E***, REC+ values were compared between groups. ***F***, Representative images of retinal histology from each of the genotypes. ***G***, Quantitation of photoreceptor nuclei number in a row in the ONL. ***H***, Representative confocal images of retinal cross-sections stained with anti-rhodopsin (green) and DAPI (magenta). Key: GCL, ganglion cell layer; INL, inner nuclear layer; ONL, outer nuclear layer; RPE, retinal pigmented epithelium. Panel widths: D, 4 mm; F, 297 μm (*Ush1g+/+*), 230 μm *(Ush1g^bw^/+*), 365 μm *(Ush1g^bw/bw^);* H, 150 μm.

### Auditory and vestibular functional phenotype

*USH1G* mutations also cause profound deafness and constant vestibular dysfunction in humans at birth^15^. The behaviors that led to the initial isolation of the *Ush1g^bw^* allele are largely similar to those seen in other mouse lines with mutations affecting the vestibular system; the allele descriptors *“circler,” “shaker,”* and *“waltzer,”* commonly used for these lines^16^, all indicate hyperactivity and stereotyped motor behavior. Because most mouse lines with vestibular dysfunction also lack auditory function, we predicted that *Ush1g^bw/bw^* mice would have disruptions in hearing and balance. Indeed, *Ush1g^bw/bw^* mice had profound hearing loss, objectively shown with ABR measurements^17^. None of the *Ush1g^bw/bw^* mice tested had an ABR response at 8, 16, or 32 kHz, while heterozygote mice had normal ABR thresholds (Fig. 3A). Likewise, *Ush1g^bw/bw^* mice had profound vestibular dysfunction, objectively shown by threshold elevation in VsEP measurements^18^. None of the *Ush1g^bw/bw^* mice tested had a VsEP response, while heterozygote mice had normal VsEP thresholds (Fig. 3B).

**Figure 3.**
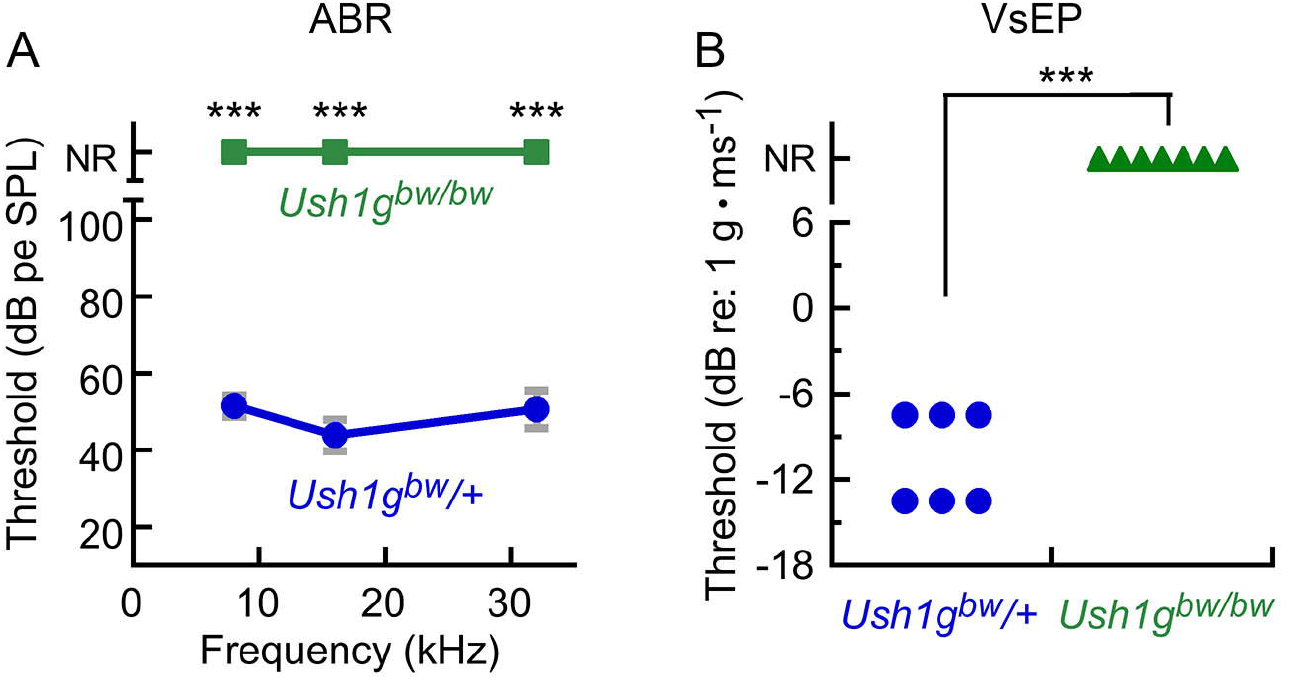
Auditory and vestibular function in *Ush1g^bw^* mice. ***A***, ABR measurements. None of the seven *Ush1g^bw/bw^* mice had an ABR response; ABRs in the seven *Ush1g^bw^*/+ control mice were normal. ***B***, VsEP measurements. None of the seven *Ush1g^bw/bw^* mice had a VsEP response; VsEPs in the seven *Ush1g^bw^*/+ control mice were normal. NR, no response. Mean ± s.e.m. plotted.

### Morphology of inner-ear hair bundles

The disruption to auditory and vestibular function and the known role of USH1G in hair cells suggested examination of their sensory hair bundles would be illuminating. Mutations in other genes that cause USH1 have severe disruption of their bundles, in large part due to loss of tip links, transient lateral links, and kinocilial links, all of which are formed by CDH23 and PCDH15 (ref. 19). Scanning electron microscopy (SEM) revealed significant disorganization of P8.5 *Ush1g^bw/bw^* bundles (Fig. 4A, E), although the disruption was not as severe as seen with homozygotes from USH1 mutant lines, including *Pcdh15^av3J/av3J^, Cdh23^v2J/v2J^*, and *Myo7a^8J/8J^* (Fig. 4B-D, F-H). Stereocilia were more coherent in *Ush1g^bw/bw^* bundles than in bundles of homozygous mice from the other strains. The most notable disruption in *Ush1g^bw/bw^* bundles was the loss of connection of the stereocilia with the kinocilium, the axonemal structure that orients a bundle. While the kinocilium degenerates after P10 (ref. 20), it still can be seen in the scanning electron micrographs of *Ush1g^KO^/+* control bundles (Fig. 4, arrows). In *Ush1g^bw/bw^* hair cells, the kinocilium was usually detached from the bundle (Fig. 4, arrows); in addition, there was a striking notch in most bundles, where the central stereocilia of each row were displaced medially (Fig. 4, arrowheads). The notch phenotype was also seen in hair cells of the other homozygous USH1 mutant lines (Fig. 4C-D, G-H), with the strongest effect seen in *Myo7a^8J/8J^* bundles (Fig. 4H).

**Figure 4.**
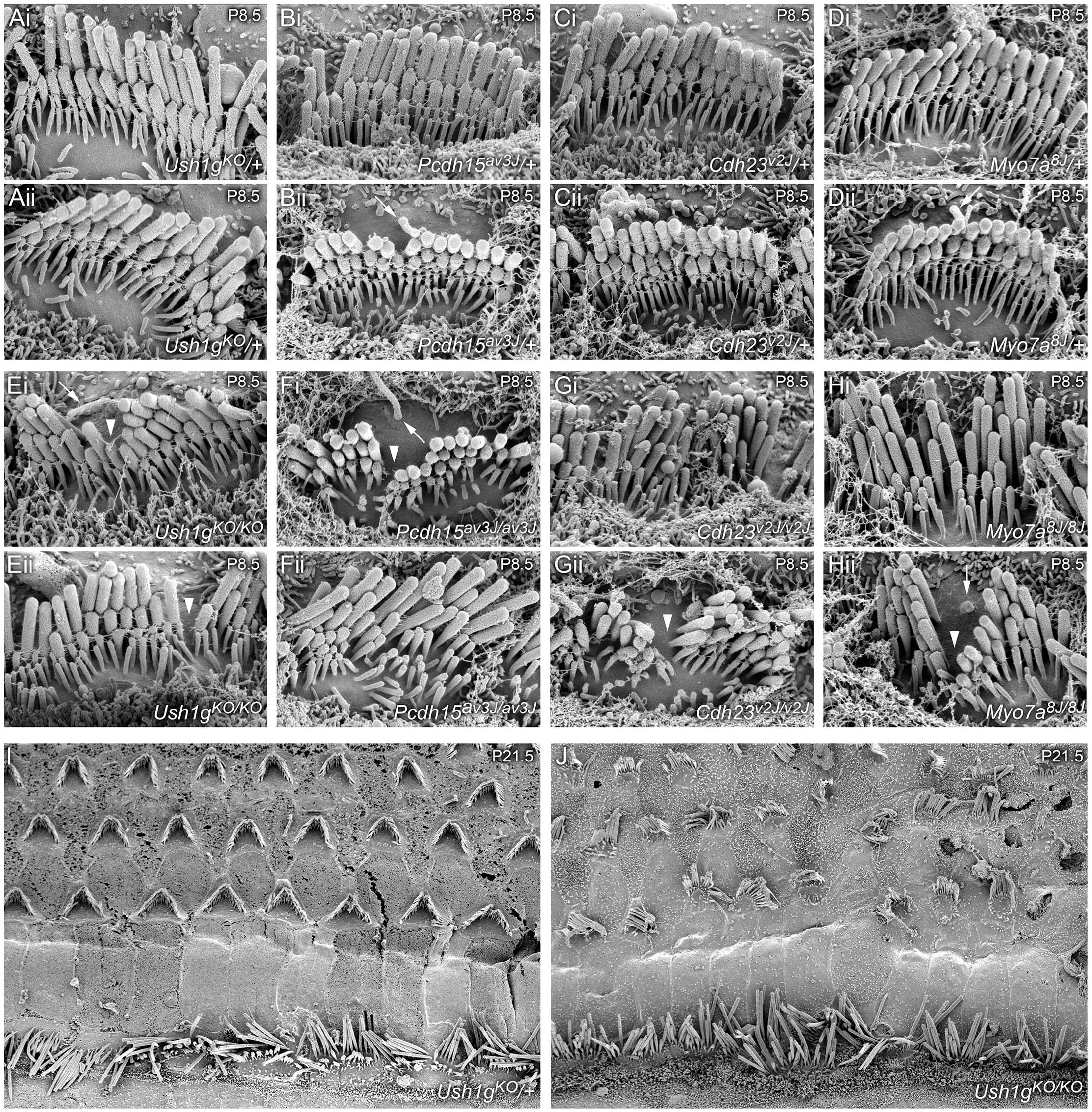
Scanning electron microscopy of hair bundles from Usher 1 mouse models. ***A-H***, Scanning electron micrographs of single IHC hair bundles from P8.5 cochleas of indicated genotypes. For each genotype, two examples (i and ii) are provided. Arrows indicate kinocilia; arrowheads indicate notch in bundles of mutant hair cells. ***I-J***, Scanning electron micrographs of P21.5 cochleas of indicated genotypes. Panel widths: A-H, 6 μm; I-J, 50 μm.

Many row 2 stereocilia in USH1 heterozygotes had beveled tips (Fig 4A-D), indicative of force production via the tip link and resulting actin remodeling^21,22^. All stereocilia tips in *Ush1g^bw/bw^* hair bundles were rounded, which suggests that functional tip links were not present (Fig. 4E); this observation was consistent with the extreme elevation of ABR thresholds. A similar rounded stereocilia tip phenotype was seen in *Pcdh15^av3J/av3J^, Cdh23^v2J/v2J^*, and *Myo7a^8J/8J^* stereocilia (Fig. 4F-H). Other links were still present in *Ush1g^bw/bw^* bundles (Fig. 4E), which may account for the increased bundle coherence compared to the other USH1 mutants (Fig. 4F-H).

At P21.5, *Ush1g^bw/bw^* OHC hair bundles were particularly disrupted compared to controls (Fig. 4I-J). Adjacent stereocilia in P21.5 *Ush1g^bw/bw^* IHC bundles were not as coordinated in length as those in *Ush1g^KO^/+* controls (Fig. 4I-J).

Using phalloidin labeling to highlight the actin cytoskeleton, we noted that utricle hair bundles were disrupted in *Ush1g^bw/bw^* mice; their stereocilia were not coherent, and projected in multiple directions (Fig. 5A). Cochlear bundles remained relatively coherent in *Ush1g^bw/bw^* mice (Fig. 5B-C), although a notch in the bundle was usually obvious (Fig. 5C, arrow).

**Figure 5.**
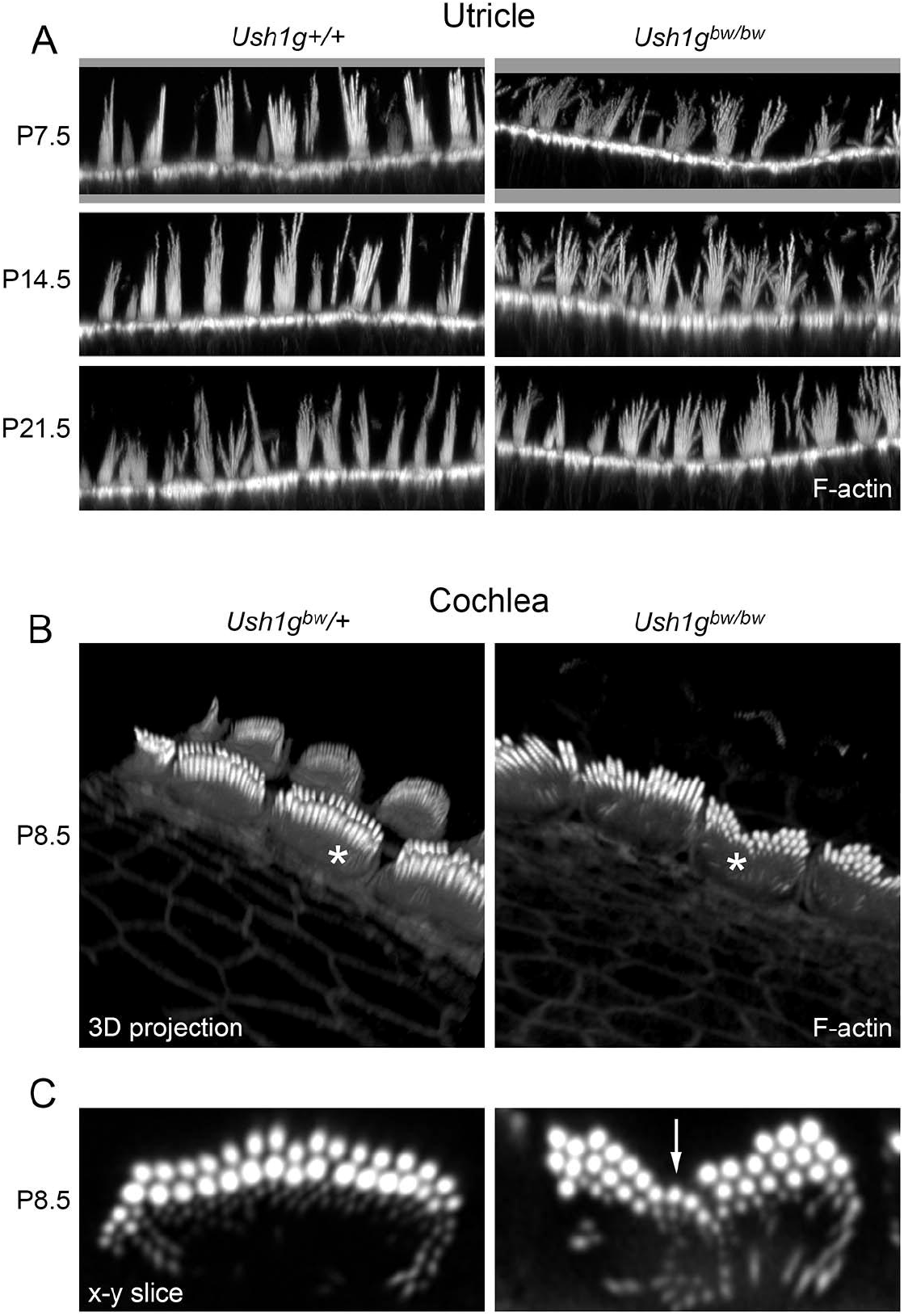
Phalloidin-labeled hair bundles from utricle and cochlea *Ush1g^bw^* hair cells. ***A***, Utricle from P7.5 through P21.5. ***B-C***, Cochlea. Panels in B are 3D projections of an x-y-z stack. Panels in C show single x-y slices through IHCs indicated with asterisks in B. Arrow in right panel of C indicates prominent notch in the hair bundle. Panel widths: A, 84 μm; C, 10 μm.

### Distribution of row-specific proteins in cochlear hair cells

In our hands, immunolocalization with an antibody reported to be specific for USH1G^23^ showed similar patterns in *Ush1g*^*bw*^/+ controls and *Ush1g^bw/bw^* mutants (not shown), i.e., it was not specific. We also used antibodies against acetylated tubulin to mark the tip of the kinocilium^24^. Similar to results seen when the kinocilium was localized using scanning electron microscopy, acetylated tubulin was associated with the central stereocilia in *Ush1g*^*bw*^/+ hair bundles (arrows in Fig. 6A-B), while it was less consistently found adjacent to stereocilia in *Ush1g^bw/bw^* bundles (Fig. 6A-B). The immunolocalization and scanning electron microscopy results indicate that USH1G is essential for function of the kinocilial links.

**Figure 6.**
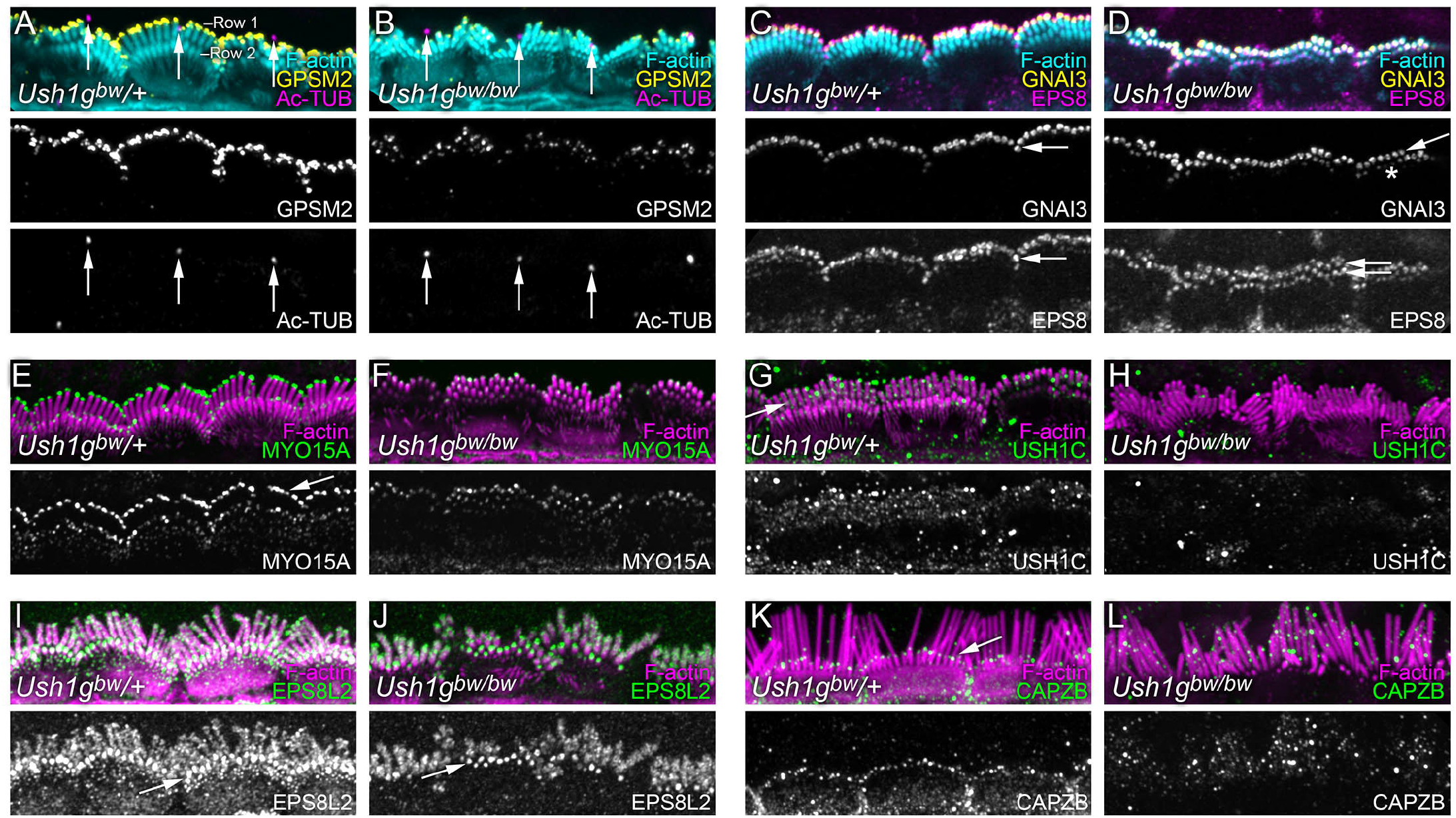
Localization of row-specific proteins and other proteins in *Ush1g^bw^* hair cells. ***A-B***, GPSM2 and acetylated tubulin (Ac-TUB). GPSM2 levels decrease in *Ush1g^bw/bw^* hair bundles but the row 1 concentration is unchanged. Arrows indicate kinocilia tips. ***C-D***, GNAI3 and EPS8. GNAI3 is largely still row 1-specific (arrows) in *Ush1g^bw/bw^* bundles but some row 2 labeling is apparent (asterisk). EPS8 shifts from highly row 1-specific in *Ush1g^bw^*/+ bundles (arrow) to equally distributed in rows 1 and 2 in *Ush1g^bw/bw^* bundles (double arrows). ***E-F***, MYO15A. MYO15A is abundant in row 1 tips in *Ush1g*^*bw*^/+ bundles (arrow) but reduced there in *Ush1g^bw/bw^* bundles. ***G-H***, Punctate USH1C labeling in *Ush1g*^*bw*^/+ bundles (arrow) is missing in *Ush1g^bw/bw^* bundles. ***I-J***, EPS8L2. EPS8L2 is normally concentrated at row 2 tips but is found throughout stereocilia membranes at low levels; this pattern is unchanged in *Ush1g^bw/bw^* bundles. ***K-L***, CAPZB. CAPZB labeling is present at row 2 tips in *Ush1g^w^/+*bundles (arrow) but is found throughout *Ush1g^bw/bw^* bundles in a punctate pattern. Panel widths, 30 μm.

A protein complex made of GPSM2, GNAI3, WHRN, MYO15A, and EPS8 concentrates at the tips of row 1 stereocilia in cochlear inner hair cells^25–27^; each of these proteins is required for normal lengthening of row 1 stereocilia^27^. We used immunocytochemistry to localize proteins of the row 1 complex in *Ush1g^bw^* IHCs (Fig. 6). GPSM2 is normally exclusively found at row 1 tips; in *Ush1g^bw/bw^* hair bundles, it was also concentrated only in row 1 (Fig. 6A). GNAI3 had a similar distribution (Fig. 6C-D), although more GNAI3 was found at tips of row 2 stereocilia in *Ush1g^bw/bw^* bundles (asterisk in Fig. 6D). By contrast, EPS8 shifted from having most labeling at row 1 tips in *Ush1g*^*bw*^/+ bundles (arrow in Fig. 6C) to nearly identical levels at both row 1 and 2 tips in *Ush1g^bw/bw^* bundles (double arrow in Fig. 6D). This broadened distribution of EPS8 resembled that seen in *Pcdh15^av3J/av3J^* and *Cdh23^v2J/v2J^* mutants^28^. MYO15A showed a similar altered distribution; while most MYO15A was found at row 1 tips in *Ush1g*^*bw*^/+ bundles (Fig. 6E), its overall level was reduced in *Ush1g^bw/bw^* bundles and it was evenly distributed between rows 1 and 2 (Fig. 6F).

USH1C (harmonin) is a binding partner for USH1G at the upper insertion of the tip link^29^. In *Ush1g*^*bw*^/+ bundles, USH1C was distributed throughout the hair bundle, but appeared to concentrate in a line on row 1 stereocilia that could correspond to the tip-link upper insertions (arrow in Fig. 6G). USH1C labeling disappeared in *Ush1g^bw/bw^* bundles (Fig. 6H), consistent with a requirement of USH1G for proper localization of USH1C.

Several proteins concentrate at tips of row 2 stereocilia in developing hair bundles, including EPS8L2, ESPNL, BAIAP2L2, CAPZB, TWF2, DSTN, and CFL1, and the locations of most of these are regulated by mechanotransduction^30–35^. We examined the location of two of these, EPS8L2 and CAPZB, in *Ush1g^bw^* hair cells. In *Ush1g*^*bw*^/+ bundles, EPS8L2 was found along the shafts of all stereocilia, but was also strongly concentrated at row 2 tips (Fig. 6I); the distribution of EPS8L2 was similar in *Ush1g^bw/bw^* bundles (Fig. 6J). By contrast, CAPZB no longer was found at row 2 tips in *Ush1g^bw/bw^* bundles (Fig. 6L), unlike its robust tip localization in row 2 of *Ush1g*^*bw*^/+ controls (Fig. 6K).

## Discussion

The *Jackson shaker* (*js*) allele of *Ush1g* was originally identified in 1967 (ref. 36). The *js* allele was mapped to the distal part of chromosome 11, allowing identification of the *Ush1g* gene^37^. Simultaneously, human Usher syndrome type 1G (USH1G) was mapped to the *USH1G* gene^14^. The protein product was termed sans (scaffold protein containing ankyrin repeats and SAM domain), although we refer it to here by the protein symbol derived from its gene symbol (USH1G).

As with other USH1 genes, including *Cdh23, Pcdh15, Ush1c*, and *Myo7a, Ush1g* is primarily expressed in the brain, testis, eye, and inner ear^37,38^. USH1G is particularly well characterized in the inner ear. There, CDH23 and PCDH15 form the hair cell’s tip links, and USH1G, USH1C, and MYO7A anchor the CDH23 end of the tip link^39^. MYO7A, which interacts with USH1C, also forms a strong complex with USH1G^40,41^; moreover, USH1G and USH1C bind tightly together^40,42^. Finally, MYO7A, USH1C, and USH1G form a liquid-phase biomolecular condensate in vitro, which suggests that these three proteins form a multivalent, stable complex^43^. Thus in the inner ear, and probably in the retina too, MYO7A, USH1C, and USH1G form a stable protein complex that can anchor CDH23 and transmit force generated by MYO7A’s motor activity.

We identified a new allele for *Ush1g*, called *Ush1g^bw^*, which occurred unexpectedly when *Neil1^-/-^ mice* developed spontaneous behavioral changes, presumably because of the lack of base-excision repair with the loss of NEIL1 protein. The *Ush1g^bw^* allele may be useful for dissection of auditory and vestibular function; this allele replicates the constellation of behavioral and morphological phenotypes seen in mouse lines with disruptive mutations in other USH1 genes, including *Cdh23, Pcdh15*, and *Myo7a*. Results with the new allele confirmed the essential role USH1G plays in establishing and operating key links within the hair bundle, including tip links, transient lateral links, and kinocilial links.

### Backwards walking phenotype

Backwards walking has not been reported as a phenotype for other *Jackson shaker* alleles of *Ush1g* (https://www.informatics.jax.org/marker/MGI:2450757). The original *js* allele has a frameshift mutation that leads to a truncated protein at amino acid 245, which deletes the C-terminal SAM domain but leaves intact the ankyrin repeats, sterile α-motif (SAM) domain, and 95-aa central region. The *js-2J, js-3J*, and *js-seal* alleles all introduce mutations that may produce truncated proteins but not fully eliminate expression. Given polypeptide chain termination following the third amino acid, *Ush1g^bw^* is surely a true null; consequentially, it may have more disruptive phenotypes compared to the other *Jackson shaker* alleles. Indeed, it is not uncommon for phenotypes of mouse alleles with naturally occurring mutations to be less profound than those of nulls, so the *Ush1g^bw^* mouse line may be particularly useful for further characterization of auditory and vestibular function.

Backwards walking has also been reported as a phenotype in other auditory and vestibular mutant mouse lines, including *tailchaser (Myo6*)^44^ and *spinner (Tmie*)^45^. Whether backwards walking also is present in other mutant lines that show hyperactivity, circling, and other connected vestibular phenotypes is not clear. All evidence suggests that this phenotype is due to auditory and vestibular disruption, however, not due to a new role for USH1G in other tissues.

### Retinal phenotype

Mouse models of USH1 are congenitally deaf but show minimal signs of retinal degeneration^46–48^. Although some models showed modest reductions in ERG, these changes could either be due to a phenotype resulting from the loss of an USH1 protein or be a result of variation^46^. Our model was consistent with previous USH1 models in that there was no measurable retinal degeneration in *Ush1g^bw/bw^* mice by P180. The USH1 proteins co-localize at the calyceal processes in the retina, which is a membrane-membrane connection site between a photoreceptor’s outer segment and its inner segment. Mouse photoreceptors lack calyceal processes, however, and USH1 proteins cannot be detected in the outer-segment basal region^47^. Without the proper cellular structure, mouse models cannot recapitulate the retinal phenotype associated with USH1. To overcome this limitation, CRISPR/Cas9 editing of the *MYO7A* gene has recently been used to generate an *USH1B* rhesus macaque model^49^, while the *USH1C* gene was edited to produce an *USH1C* pig model^50^; each model will be useful in characterizing retinal degeneration.

### Hair-cell phenotype

USH1G interacts with both CDH23 and PCDH15^23^, and we found that the morphological phenotype of *Ush1g^bw^* strongly resembled those of *Cdh23^v2J^* and *Pcdh15^av3J^*. One of the more striking alterations seen in hair bundles from these three mutant lines was the loss of attachment of the kinocilium to the bundle and a resulting notch in the bundle. As CDH23-PCDH15 kinocilial links anchor the tallest stereocilia to the kinocilium^51,52^, these results suggest that USH1G is an essential component of the intracellular anchors for these links. USH1G is well known for its critical role in forming the motor complex at the bundle’s upper tip-link density^23^, which controls tension in tip links; while the bundle fragmentation and kinocilial displacement has been noted previously in outer hair cells using a late-knockout *Ush1g* model^23^, a role of USH1G role in kinocilia links themselves has not been previously proposed.

As previously noted^19^, USH1C labeling is absent in hair bundles from mice with disruptions in *Ush1g*.Given that USH1C interacts strongly with USH1G^40,42^, we agree that USH1C is incapable of being delivered to stereocilia in the absence of USH1G^53^. The USH1G PDZ binding motif (PBM) and SAM domain form a highly stable complex (K_D_ ≈ 2 nM) with the combined N-terminus/PDZ1 domain (NPDZ1) of USH1C^53^. In this tight complex, USH1C is still capable of binding to CDH23^53^. USH1G is also essential for tight interaction of USH1C with MYO7A^53^.

Interaction of USH1G with PCDH15 is less well understood. Double heterozygous *Pcdh15^av3J^/+;Ush1g^js^*/+ mice showed elevated ABR thresholds, however, demonstrating a genetic interaction that may reflect a physical association^54^. In support of a direct USH1G-PCDH15 interaction, tagged USH1G was recruited to the plasma membrane of COS-7 cells when PCDH15 was co-expressed but not when it was expressed on its own^23^. MYO7A interacts with PCDH15^55^, and since MYO7A and USH1G interact strongly^41^, it is likely that the functional complex of PCDH15 and USH1G also includes MYO7A. Indeed, PCDH15 is mislocalized in *Myo7a^sh1/sh1^* mutants^55^.

Homozygous null mutations in *Cdh23* and *Pcdh15* lead to altered distribution of proteins that concentrate at the tips of row 1 stereocilia in postnatal hair bundle development^28^. While localization of GPSM2 and GNAI3 was only modestly affected, EPS8 shifted from being primarily found at row 1 tips to being equally distributed between row 1 and 2 tips. We also showed that CDH23 and PCDH15 play an underappreciated role in coordinating the lengths of adjacent stereocilia^28^. Row 1 proteins showed similar altered distribution in *Ush1g^bw/bw^*, and adjacent stereocilia lengths were less coordinated, albeit not as profoundly disrupted as in *Cdh23^v2J/v2J^* and *Pcdh15^av3J/av3J^* hair cells. The protein redistribution seen in *Ush1g^bw/bw^* hair cells highlights the importance of CDH23-PCDH15 links and their intracellular partners—like USH1G—in controlling the location of proteins important for control of actin polymerization, like the row 1 complex.

Distribution of the row 2 protein EPS8L2 was relatively unaffected in *Ush1g^bw/bw^, Cdh23^v2J/v2J^*, and *Pcdh15^av3J/av3J^* hair cells, but the concentration of CAPZB at row 2 tips was eliminated in all three mutant lines. Because functional transduction is required for CAPZB localization at row 2 tips^56^, these results are consistent with there being no mechanotransduction in *Ush1g^bw/bw^* hair cells, as also suggested by the profound elevation of both ABR and VsEP thresholds.

## Conclusions

The *Ush1g^bw^* mouse line is a null mutation of *Ush1g*, which will be of utility for characterizing USH1G in tissues that express the protein, including the retina, cochlea, and vestibular system. Alterations in hair-bundle structure in *Ush1g^bw/bw^* mice suggest that USH1G contributes to anchoring kinocilia links, not just tip links. In cochlear hair cells, the *Ush1g^bw/bw^* phenotype is similar to those seen in *Cdh23^v2J/v2J^* and *Pcdh15^av3J/av3J^* mice, suggesting that USH1G is a necessary subunit in complexes with these two large cadherins.

## Abbreviations

ABR: auditory brainstem response
GCL: ganglion cell layer
INL: inner nuclear layer
ONL: outer nuclear layer
RPE: retinal pigmented epithelium
s.e.m.: standard error of the mean
SEM: scanning electron microscopy
USH1: Usher I
VsEP: vestibular evoked potential.

## Acknowledgements

We carried out confocal microscopy in the OHSU Advanced Light Microscopy Core @ The Jungers Center and electron microscopy in the OHSU Multiscale Microscopy Core. PGBG was supported by National Institutes of Health grant R01DC002368. RSL was supported by National Institute of Environmental Health Sciences grant R01ES031086, as well as the Oregon Institute of Occupational Health Sciences at Oregon Health & Science University (via funds from the Division of Consumer and Business Services of the State of Oregon; ORS 656.630). RR was supported by Casey Eye Institute Core Grant P30 EY010572 from the National Institutes of Health and unrestricted departmental funding from Research to Prevent Blindness.

## Supplemental Video

Supplemental Video S1. **Backwards-walking and hyperactivity phenotypes in *Ush1g^bw^* mice**.

